# Genome Assemblies of the Warthog and Kenyan Domestic Pig Provide Insights into Suidae Evolution and Candidate Genes for African Swine Fever Tolerance

**DOI:** 10.1101/2021.12.17.473133

**Authors:** Wen Feng, Lei Zhou, Pengju Zhao, Heng Du, Chenguang Diao, Yu Zhang, Zhen Liu, Wenjiao Jin, Jian Yu, Jianlin Han, Edward Okoth, Raphael Morode, Jian-Feng Liu

**Affiliations:** National Engineering Laboratory for Animal Breeding; Key Laboratory of Animal Genetics, Breeding and Reproduction, Ministry of Agriculture; College of Animal Science and Technology, China Agricultural University, Beijing, 100193, China; Shenzhen Kingsino Technology Co., Ltd., Shenzhen, 518107, China; Hainan Institute, Zhejiang University, Yongyou Industry Park, Yazhou Bay Sci-Tech City, Sanya 572000, China; International Livestock Research Institute (ILRI), Nairobi, 00100, Kenya; CAAS-ILRI Joint Laboratory on Livestock and Forage Genetic Resources, Institute of Animal Science, Chinese Academy of Agriculture Sciences (CAAS), Beijing, 100193, China

**Keywords:** Warthog, Kenyan domestic pig, genome assembly, African swine fever, lactate dehydrogenase B

## Abstract

As warthog (*Phacochoerus africanus*) has innate immunity against African swine fever (ASF), it is critical to understand the evolutionary novelty of warthog to explain its specific ASF resistance. Here, we present two completed new genomes of one warthog and one Kenyan domestic pig, as the fundamental genomic references to decode the genetic mechanism on ASF tolerance. Our results indicated, multiple genomic variations, including gene losses, independent contraction and expansion of specific gene families, likely moulded warthog’s genome to adapt the environment. Importantly, the analysis of the presence and absence of genomic sequences revealed that, the warthog genome had a DNA sequence absence of the lactate dehydrogenase B *(LDHB)* gene on chromosome 2 compared to the reference genome. The overexpression and siRNA of *LDHB* indicated that its inhibition on the replication of ASFV. Combining with large-scale sequencing data of 123 pigs from all over the world, contraction and expansion of *TRIM* genes families revealed that *TRIM* family genes in the warthog genome were potentially responsible for its tolerance to ASF. Our results will help further improve the understanding of genetic resistance ASF in pigs.

## Introduction

Suidae is a family of artiodactyl mammals that originated 20 to 30 million years ago (mya) and consists of 15 to 17 extant species that are grouped into five genera (Frantz et al. 2016). As the most abundant and widely distributed member of the Suidae family, the domestic pig (*Sus scrofa domesticus*) is a domesticated species of global importance, as it constitutes the preferred source of animal protein for human consumption and serves as an important biomedical model. The common warthog (*Phacochoerus africanus*) is a wild member of the Suidae family that is naturally distributed in the grasslands, savannas, and woodlands of sub-Saharan Africa. The large and flat head of warthog is covered with protective bumps and armed with four sharp tusks, presenting a much different appearance from that of domestic pig.

Adaptive evolution driven by natural selection plays a fundamental role in species diversification. The unique geographical environment provides the Suidae with an African genetic basis in relation to their phenotypic traits, including pathogen challenge and fitness in resource-limited systems. For instance, warthogs have been shown to have natural immunity against African swine fever (ASF), and has no clinical symptom of the disease (Heuschele and Coggins 1969). ASF is mainly transmitted among the wildlife hosts of the African swine fever virus (ASFV), and it is a highly communicable and fatal infectious disease in the swine industry, with a fatality rate of 100%. ASFV can be spread both through wild boars and domestic pigs. The ASF has caused serious damage to the global pig industry, and it continuously threats the swine industries of many African and Asian countries.

Over the past decade, many genome assemblies have been assembled and released to investigate the evolution, speciation, and introgression of the Eurasian Suidae. The current reference genome *Sus scrofa* 11.1 has an N50 scaffold length of 48.2 Mb and a total sequence length of 2.5Gb in 706 scaffolds (Warr et al. 2020). However, most of the studies have only focused on Eurasian domestic pigs, and there is a lack of a suitable reference genome or sufficient re-sequencing data for the African Suidae. Therefore, there is a huge knowledge gap about the African Suidae genome, and it limits our understanding of the evolution of Suidae. For instance, the breed composition and genetic diversity of African domestic pigs, especially in East Africa, have not been fully understood. The historical events leading to the formation of the East African pig lineages remains poorly explored because of the lack of sufficient genetic and archaeological evidence. In addition, the identification of novel genetic variants and gene flows among African domestic pigs may help us provide more reasonable answers to numerous genetic and historical research questions, including possible independent domestication of African pigs and the impact of European colonization on the African pigs’ domestication.

In the present study, we generated two *de novo* genome assemblies of the African Suidae with one warthog and one Kenyan domestic pig, which could enhance our understanding of the evolution of African Suidae and unravel of the molecular basis of their unique phenotypes. The fully annotated genomes allowed us to identify the genetic content of presence/absence variation (PAV) in African Suidae, and provided us new insights into Suidae genome evolution. We presented the speciation history of the African domestic pigs, providing the evidence in support of gene flow between Kenyan and European domestic pigs through the population genetic analysis of whole-genome re-sequencing data. Besides, the orthologous genes in Suidae were used to reveal the gene change of the warthog genome responsible for their adaptation to the African environment, involving gene family expansions and contractions. The results provided us new candidate genes for understanding the molecular mechanism of warthog resistance to ASF.

## Results

### *De novo* assembly of the Kenyan domestic pig and warthog genomes

To present comprehensive genomic resources for the Suidae family in Africa, we performed two new genome assemblies for one wild common warthog and one Kenyan domestic pig using the 10× Genomics “Linked-Read” sequencing technology. We generated 1,202 million and 1,179 million 150-nt chromium-linked paired-end reads, which resulted in a genome coverage depth of ∼69× (180.32 Gb) and ∼68× (176.97 Gb), respectively (**Supplementary Table 1**).

The warthog and Kenyan domestic pig genomes were assembled and scaffolded using a Supernova assembler with the diploid pseudo-haplotype style, and the resulting scaffolds were gap-filled using the Sealer application (Paulino et al. 2015). After removing the contamination from adaptors and viral genomes, the final assemblies yielded 19,366 scaffolds in a total length of 2.417 Gb with an N50 of 13.75 Mb and 13,380 scaffolds in a total length of 2.445 Gb with an N50 of 30.52 Mb for the warthog and Kenyan domestic pig genomes (**Supplementary Table 2**), respectively.

The quality of our genome assemblies was evaluated (**Supplementary Table 3**) with the following steps. First, we mapped all chromium-linked paired-end reads to our genome assemblies with no large structural variation and an average sequence identity of 99.32%, which indicated that our assemblies were correct and contained almost all of the information in the raw reads. A further assessment of genome completeness revealed that our genome assemblies recovered 91.5% and 93.0% of the 4104 single-copy orthologs in mammalian gene groups from the warthog and Kenyan domestic pig, respectively, which were comparable to the BUSCO completeness score of 93.0% for the reference genome build from a Duroc pig (*Sus scrofa* 11.1). We observed that 89.6% and 96.26% of 41,909 transcripts from the pig reference genome could be mapped to the warthog and Kenyan domestic pig assemblies (≥90% query coverage), respectively. Altogether, these results indicated the high quality of our *de novo* assembled genomes of warthog and Kenyan domestic pig.

### Gene prediction and genomic TE composition

Using 227,840 orthologous protein isoforms (*Sus scrofa* species) from OrthoDB 10.1 (https://www.orthodb.org/) as gene evidence, we employed the BRAKER2 pipeline to predict protein-coding genes (PCGs) by integrating the *ab initio* and homology-based approaches.

As a result, 17,781 and 22,127 PCGs were identified in the warthog (with an average of gene length in 21,907 bp and 6.7 exons per gene) and Kenyan domestic pig (21,929 bp and 6.5 exons) assemblies, of which 59.3% and 52.4% were homologous with the 27,445 PCGs annotated in the pig reference sequence, respectively. We found 1,668 novel PCGs (**Supplementary Table 4**) in the warthog assembly, which were supported by mRNA transcripts and had an ortholog in either the non-redundant database (NR) of NCBI, InterPro, or UniportDB (Swiss-Prot and TrEMBL). Notably, these warthog unique PCGs contained the largest percentage of immune-related genes as compared to pig PCGs and were significantly enriched in KEGG pathways related to ‘‘Pathways in cancer’’ and ‘‘Metabolic pathways’’ (adjusted p-value < 0.01), which could explain why the warthogs are very strong in genetic resistance/tolerance to a variety of endemic parasitic and viral diseases.

Identifying repetitive sequences can effectively improve the accuracy of genome annotation. Transposable elements (TEs), one of the main and ubiquitous genomic repetitive elements (**Supplementary Table 5**), are the driving forces in shaping genomic architecture and evolution. In total, 4,039,334 TEs occupied 42.82% (1,053 Mb) of the entire warthog genome, two-thirds of them were assigned to a specific family. In total, 4,247,450 TEs occupied 42.94% (1,061 Mb) of the entire Kenyan domestic pig genome. As observed in previous studies (Santos et al. 2014; Chen et al. 2019), the most abundant TEs were retrotransposons (accounted for ∼95% of TEs). Both LINEs and SINES together accounted for 82% and 81%, while DNA transposons took only 5.5% and 5.7% of the warthog and Kenyan domestic pig genomes (**Fig. 1a**). LINEs accounted for more than half of TEs, thereby being the most abundant pig TEs in the both two assembled genomes, which were contradictory to the early observation that more quantity of SINEs (66.8%)than LINEs (14.1%) in Chinese and European pigs (Zhao et al. 2016). Family L1/CIN4 had major contributions, accounting for 93.3% and 92.6% of classified LINEs in the warthog and Kenyan domestic pig genomes, respectively. Then Kimura distance-based copy divergence analysis and correction with the CpG content of each TE were performed using RepeatMasker. The divergence pattern indicated the most ancient SINEs at 20 mya and one burst of LINEs at 60 mya, but no recent expansion of TEs in the two genomes. The warthog and Kenyan domestic pig genomes shared a similar pattern expansion and contraction of Tes (**Fig. 1b**).

**Fig. 1.**
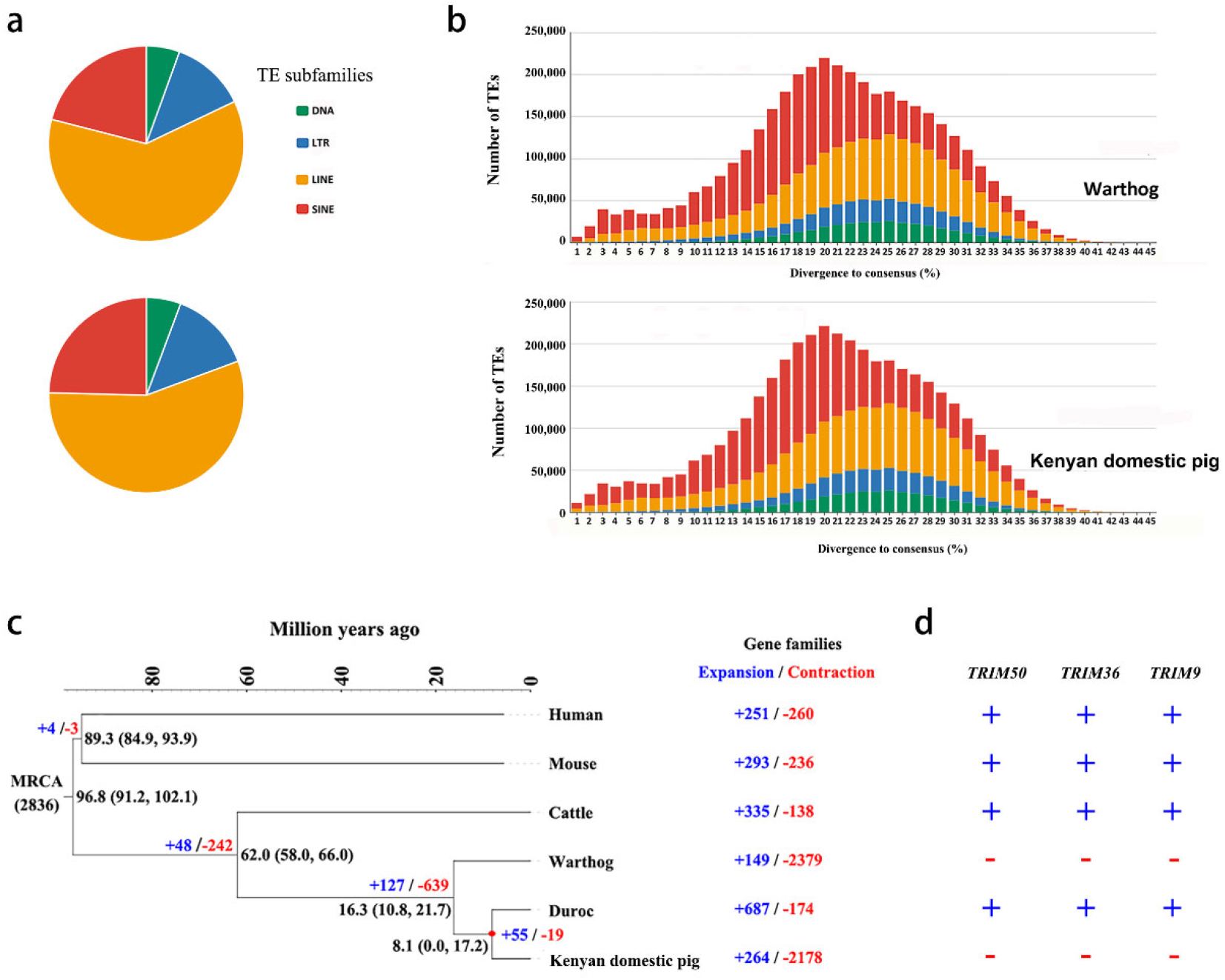
Genomic structure of the warthog and Kenyan domestic pig genomes and gene prediction. **a**. Proportions of DNA transposons, LTR, LINE, and SINE retrotransposons in the two assembled genomes. **b**. History of TE accumulation in the warthog and Kenyan domestic pig genomes. The stacking plots were used to show the divergence distribution either by summing all four TE classes or separately for each family. **c**. A phylogenetic tree was constructed with 423 orthologs from six mammals using OrthoMCL with a Markov cluster algorithm. Divergence time was estimated with the approximate likelihood calculation method in conjunction with a molecular clock model. A bar within a branch indicates the 95% confidence interval of divergent time. The positive and negative numbers adjacent to the taxon names are gene family numbers of expansion/contraction obtained from the CAFE analysis. **d**. The expansion or contraction of three TRIM family genes (TRIM50, TRIM36 and TRIM9). The “+” means expansion and “-” means contraction.

### The expansion and contraction of gene family evolution

A computational analysis of gene family evolution (CAFE) revealed a huge contraction of orthologous gene families in the warthog genome based on the observation of 2,379 contracted but 149 expanded gene families (**Fig. 1c and Supplementary Tables 6 and 7**). Among these expanded gene families, 104 families were unique to the warthog. Massive expansions of phospholipid-transporting ATPases, which have been previously noted in the pig reference genome, have functional implications for muscle electrical conductivity (Mármol-Sánchez et al. 2020) and thermotolerance (Kim et al. 2018). The expanded gene *NFKB1* identified in the warthog plays a crucial role in the NF-KB pathway (Fliegauf et al. 2015). In the Kenyan domestic pig genome, 2,178 gene families were contracted while 264 were expanded. Among the 37 expanded gene families shared by the warthog and Kenyan domestic pig, genes *MX1, MX2, ETS1*, and *PSPC1* were involved in immune response (Wang et al. 2017; Lee et al. 2019; Fan et al. 2020). Interestingly, we found three contracted TRIM genes (*TRIM 50, TRIM36*, and *TRIM9*) in both warthog and Kenya domestic pig but they were expanded in the Duroc pig (**Fig. 1d**).

### Massive presence-absence sequences among the Suidae genomes

Intraspecific presence/absence variants (PAVs) are important sources of genetic diversity and divergence that contribute to an organism’s ability to adapt to its specific habitat. To capture the set of PAVs in all available accessions of Suidae, we performed a multi-genome alignment of 16 genomes to the pig reference genome and retained the unaligned scaffolds and the structure variations (insertion or deletion longer than 50 bp) from aligned scaffolds as the PAVs. As expected, compared with all pig breeds, the warthog had the highest number of PAVs (165,452), ranging from 31,939 in 13.3 Mb to 98,767 in 27.8 Mb (**Supplementary Table 8**). A further exploration of genomic distribution revealed that the density of PAVs decreased from distal region(s) toward the centromere within a chromosome and showed a significant enrichment in the intergenic regions (chi-squared test, P < 0.05).

By comparing the distribution patterns of PAVs and protein coding regions along the genomes, we found 67 protein coding regions to be linked with PAVs in the reference pig genome, 19 of them had known annotations (**Supplementary Table 9**). The distribution of PAVs linked with these 19 genes in different species/breeds are shown in **Fig. 2a**. Interestingly, the *LDHB* gene was mapped to two distinct genomic locations of the pig reference genome: chromosome 5 (chr5: 51810932–51831218) and (chr2: 12988754–12990033). The entire sequence of *LDHB* that mapped to chromosome 2 of *Sus scrofa11*.*1* was missing in the warthog (**Fig. 2b**). In the meanwhile, only the copy of the *LDHB* gene on chromosome 5 was present in the warthog genome.

**Fig. 2.**
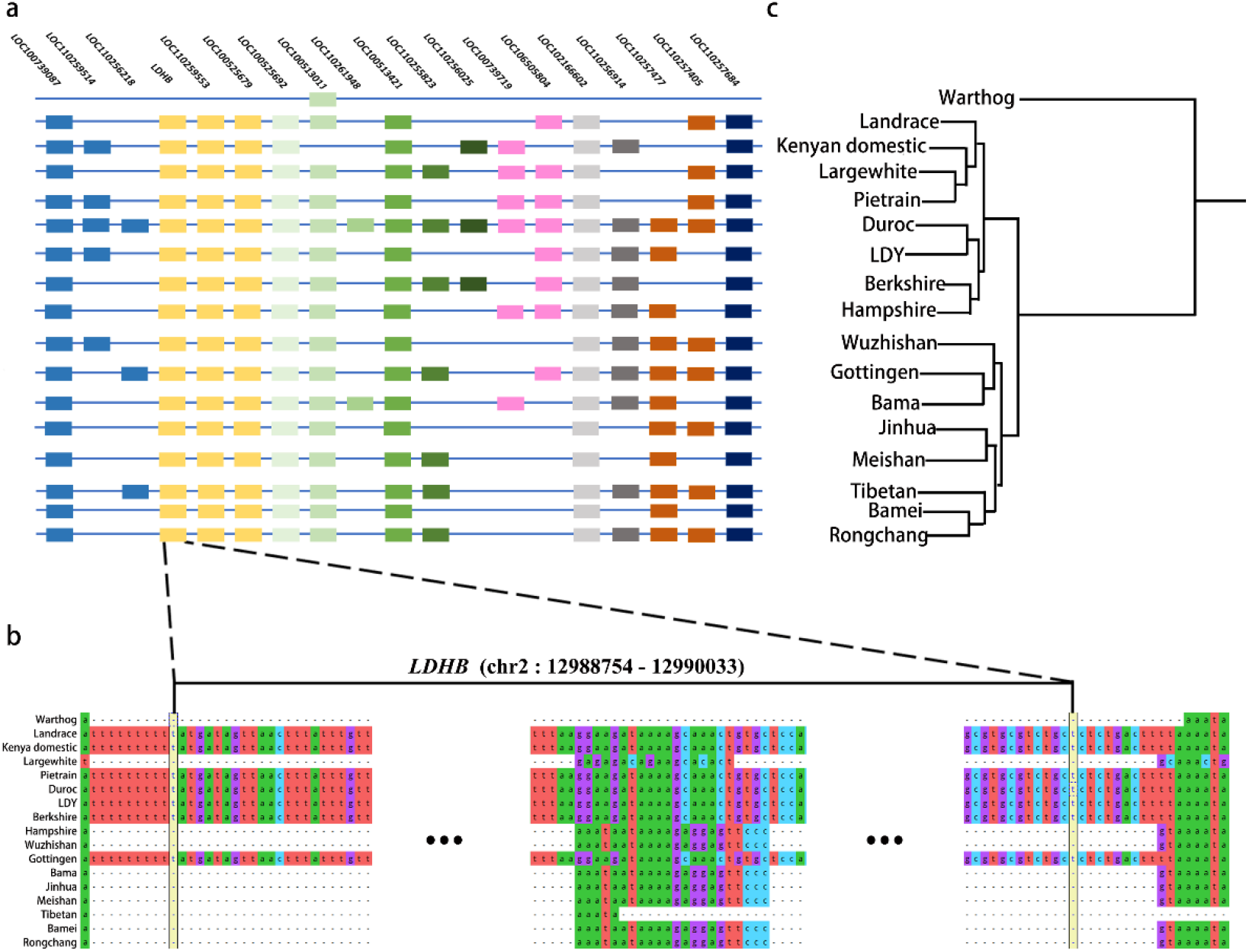
The distribution of presence and absence variation (PAV) in different pig populations. a. The distribution of PAVs that affect the coding regions of 19 genes with known annotations. The same color indicates that these genes are located on the same chromosome. b. Coding sequence alignment of gene *LDHB* in chromosome 2 of the 17 assembled genomes. c. Demographic history of pigs using 17 assembled genomes. The genomes of warthogs and Kenyan domestic pigs were assembled in this study, while 15 additional genomes were downloaded from public databases.

The expression levels of the *LDHB* gene in the heart, lung, liver, and spleen of warthogs were compared with those of Kenyan domestic pigs. The expression level of the *LDHB* gene in the four tissues was significantly higher in warthogs than in Kenyan domestic pigs by more than 10 times (**Fig. 3a)**.

**Fig. 3.**
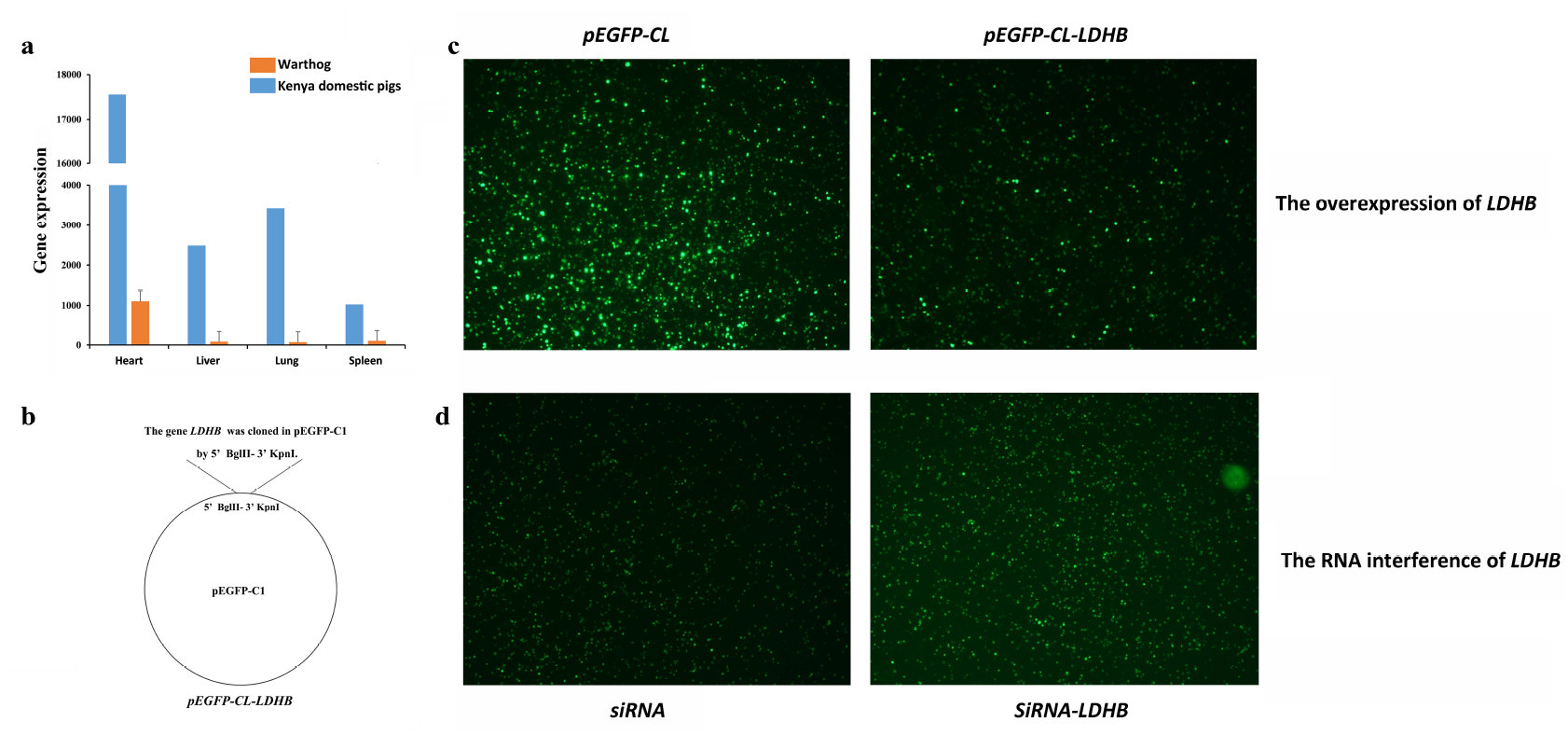
Validation of the effect of *LDHB* against ASFV. a. Gene expression of *LDHB* in the heart, liver, lung, and spleen of warthogs and Kenyan domestic pigs. b. The gene *LDHB* has been cloned intopEGFP-C1 by BglII-KpnI. c. The expression of fluorescent proteins in plasmid-transfected 3D4/21 cells after being infected with ASFV SY18 strain when *LDHB* was overexpressed. d. The expression of fluorescent proteins in plasmid-transfected 3D4/21 cells after being infected with ASFV SY18 strain when the RNA of *LDHB* was interfered with. The stronger the fluorescence intensity, the greater the amount of ASFV protein.

To determine the replication efficiency of ASFV treated *in vitro* with the overexpression and siRNA of *LDHB* gene, the 3D4/21 cells being transfected by either pEGFP-Cl-LDHB or siRNA-LDHB vector were infected with the ASFV SY18 strain (MOI = 5). The gene LDHB has been cloned into pEGFP-C1 by BglII-KpnI, (**Fig. 3b**) which represented the overexpression of *LDHB* could be carried out next. After 48h of the infection, the fluorescence intensity and proportion of 3D4/21 cells transfected by pEGFP-Cl-LDHB reduced compared with that of pEGFP-Cl (blank control) vector (**Fig. 3c**), which indicates the replication of ASFV was decreased when LDHB overexpressed. In contrast, the fluorescence intensity and proportion of 3D4/21 cells transfected with the siRNA-LDHB increased compared with the blank control (**Fig. 3d**). The fluorescence intensity reflected the inhibitory effect of *LDHB* on the ASFV replication.

### Phylogenetic analysis and effective population size estimation

To explore the relationship between African and Eurasian pig breeds, we combined the sequence information of these two assembled genomes with the publicly available assembled genomes of 15 domestic pig breeds (**Table 1**) for the genetic distance analysis. The distances between multiple genomes evaluated with MinHash were shown in **Fig. 2c**. As it was classified into a separate genus from all the domestic pig breeds, the warthog deeply diverged from all pig breeds while the genetic distances between different pig breeds were much close to each other. Meanwhile, a clear genetic separation existed between European and Asian pig breeds. In agree with our expectation, African domestic pigs were genetically closer to European domestic pig breeds than Asian domestic pig breeds, which was consistent with previous results (Gongora et al. 2011; Amills et al. 2013).

**Table 1.** Assemblies genomes used in the study.

| **Breeds** | **Project NO.** | **Breeds** | **Project NO.** |
| --- | --- | --- | --- |
| **Warthog** | PRJNA691462 | **Duroc (Sus scrofa 11.1)** | PRJNA13421 |
| **Kenya domestic** | PRJNA691462 | **½ Landrace- ¼ Duroc- ¼ Yorkshire (LDY)** | PRJNA392765 |
| **Bama** | PRJNA478804 | **Ellegaard Gottingen minipig** | PRJNA176189 |
| **Wuzhishan** | PRJNA144099 | **Tibetan** | PRJNA186497 |
| **Jinhua** | PRJNA309108 | **Hampshire** | PRJNA309108 |
| **Bamei** | PRJNA309108 | **Landrace** | PRJNA309108 |
| **Meishan** | PRJNA309108 | **LargeWhite** | PRJNA309108 |
| **Pietrain** | PRJNA309108 | **Berkshire** | PRJNA309108 |
| **Rongchang** | PRJNA309108 |  |  |

To explore the timing and mode of cephalopod evolution, phylogenetic relationship was reconstructed for 6,418 single-copy orthologs from six mammalian genomes using OrthoMCL (Fig. 1c), confirming the divergence between warthog and *Sus scrofa* at the Miocene (approximately 16.3 mya) (Frantz et al. 2016). To better understand the evolutionary history of the warthog and Kenyan domestic pigs, PSMC was used to estimate their effective population sizes (Ne) (**Fig. 4a**), which were combined with the corresponding historical events. Six populations, including warthog, Kenyan domestic pigs, Landrace, Duroc, and Meishan pigs, and Southern Chinese wild boars, were used for estimating their Ne. Between 2 mya and 1 mya, the number of warthogs experienced a sharp reduction. Approximately 2 mya, a large number of volcanoes continued to erupt in Kenya and caused severe and long droughts in the region (Maslin et al. 2015), which may be one of the reasons for the decline in the number of warthog populations. Approximately 200,000 years ago, the global temperature experienced drastic fluctuations with rapid changes from cold to warm, this sudden climate changes led to a significant increase in the number of wild pig species worldwide (deMenocal 2011). The current study showed that approximately 3,000 years ago, the effective population sizes of European and African domestic pigs tended to be highly similar.

**Fig. 4.**
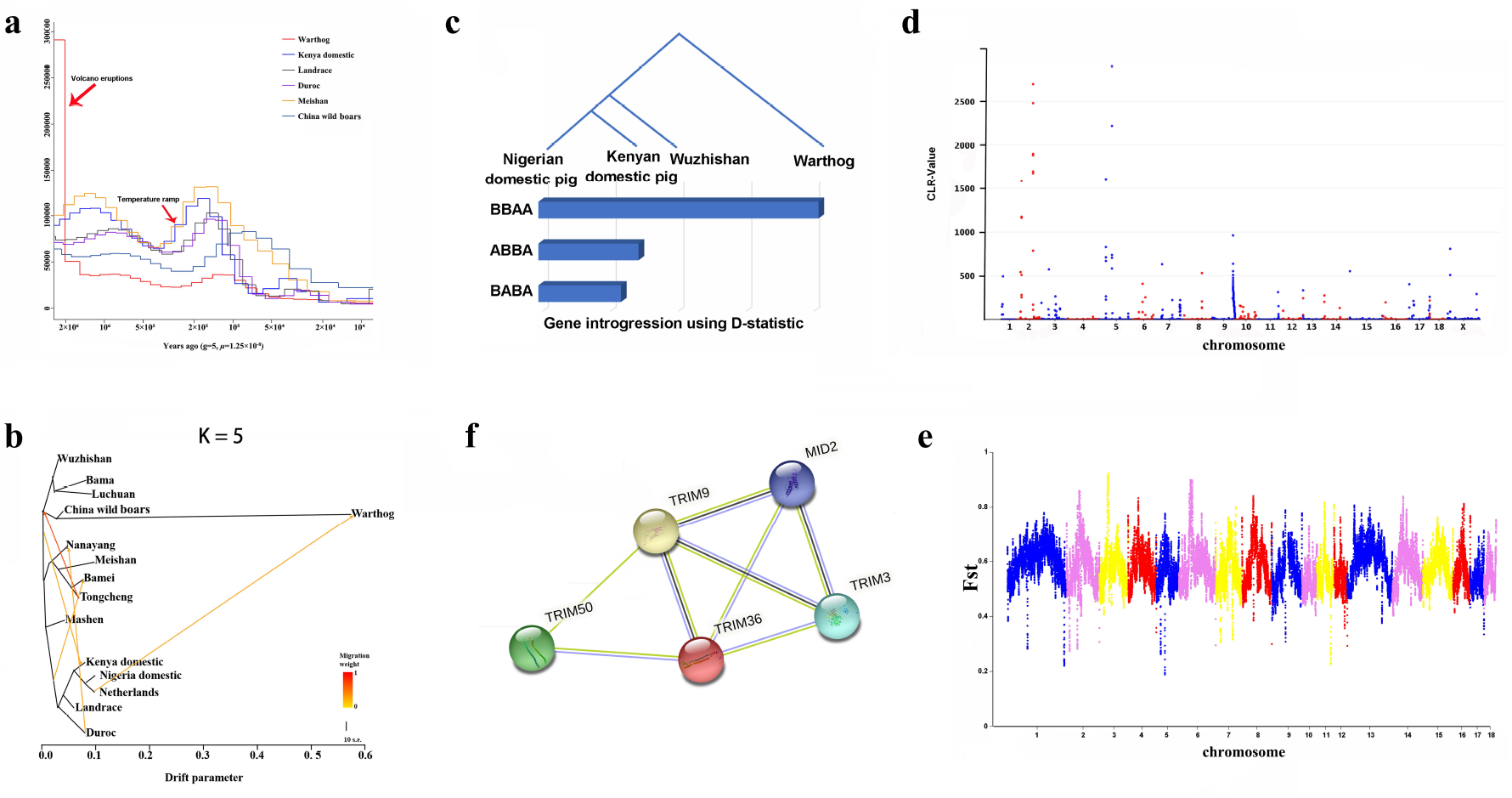
Population genetics analysis. a. Demographic history of pigs. Generation time (g) = 5; Transversion mutation rate (μ) = 1.25*10^−8^. b. Gene immigration between pig populations using TreeMix. The gene flow of the left one was 3 and the right one was 5. Red arrows indicate the gene infiltration. c. The gene introgression analysis using D-statistic. The D-statistic = 0.0962893, Z-score = 8.9732. d. Plot of CLR values in the warthog. e. Plot of Fst in the warthog and Southern Chinese wild boars. f. Protein–protein interaction (PPI) predicted networks of TRIM family genes (confidence 0.15). Each node represents one protein.

### Analysis of genetic introgression

To understand the evolutionary position of African pigs among domestic pig breeds, 123 re-sequenced whole-genomes of 14 warthog, 15 Eurasian wild boars, and 94 African and Eurasian domestic pigs from 25 breeds/populations (**Supplementary Tables 10**) were used to detect SNPs for population structure analysis, which including the phylogenetic tree (**Supplementary** (**Supplementary Figure 1**), principal component analysis (**Supplementary Figure 2**), and admixture analysis (**Supplementary Figure 3**). Combined with these population structure results, pig populations of warthogs, Kenyan domestic pigs, Nigerian domestic pigs, Netherlands wild boars, Duroc, Landrace, Mashen, Bamei, Nanyang, Meishan, Tongcheng, Bama pigs, Luchuan, Wuzhishan, and Chinese southern wild boars with at least 5 pigs each breed were analyzed for potential historical gene migration events, and the gene flow was set at 3 to 6 (**Fig. 4b**). Interestingly, when the gene flow was 3 and 6, there was a significant gene infiltration phenomenon between the warthog and the Chinese wild boar; when the gene flow was 4 and 5, there was a tendency for the Netherlands wild boar to infiltrate the warthog. No other studies on similar situations have been reported so far, and further analysis and verification are required. Gene infiltration of southern Chinese pigs into Kenyan domestic pigs was observed when the gene flow was 5 and 6. Further to verify the results, Dsuite (Malinsky et al. 2021) were employed to analyze the gene introgression when the D- statistic ≠ 0 and Z-score > 3. From D statistic results (**Supplementary Table S11**), we can easily see the gene introgressions between Afircan domestic pigs and Chinese domestic pigs. Kenya is located in East Africa, which is the crossroad of Indian Ocean trade. Studies have shown that pigs in East Africa originated in three regions: India, the Far East, and Europe (Noce et al. 2015). The results of this study demonstrated the possibility that Kenyan domestic pigs may have Asian ancestry.

### Signature analysis of selection

To test the genomic signal formed by the positive selection of the warthog in the natural environment, a selective sweep of the warthog genome was carried out using CLR. Regions with CLR-values > 100 (top 0.2%) were considered to be strongly selected (**Fig. 4c**). A total of 138 important windows were screened. Chromosomes 2 and 5 were subject to strong selection. A total of 186 genes were identified in these windows, of which 43 genes had detailed annotation information, including many interleukin family genes such as *IL19, IL1A, IL20*, and *IL24*. KEGG pathway analysis was performed on these genes, and 15 significant pathways were detected (Kanehisa et al. 2021) (Q < 0.05, **Supplementary Table 12**). These pathways were significantly related to viral infection, immunity, skin diseases, and the retina.

Based on the results of the above population structure and gene flow, we carried Fst selection signal detection on this part of the warthogs and Chinese and the Netherlands wild boars. The Fst test results showed a large difference in gene frequency between these two populations, which was consistent with expectations. Based on the selection signal results of warthogs and Chinese wild boars (**Fig. 4d**), chromosomes 2, 3, and 6 were strongly selected. The lengths of these selected regions were approximately 2.35 Mb, 4.55 Mb, and 6.35 Mb, respectively. Gene annotation was performed on these fragments. A total of 290 genes were identified, of which 219 genes had detailed annotation information. The 804 SNPs located in the 107 genes caused non-synonymous mutations (**Supplementary Figure 4; Supplementary Table 13**). The genes *RELB, IL12RB1, JAK3*, and *CREBBP* participate in the immune response, such as NF-κB, Jak-STAT, innate immunity, and other immune pathways.

Two shared genes (*MID2, SNORA70*) were detected both in selective signature results of the warthog CLR test and warthog – China wild boars Fst test. The protein encoded by *MID2* is a member of the tripartite motif (TRIM) family. Combined with TRIM family genes found in gene expansion and contraction, protein to protein analysis were used to determine the relationship among *TRIM3, TRIM36, TRIM9, TRIM50*, and *MID2*. **Fig. 4f** showed that proteins encoded by these five genes were interconnected. These genes were found involved in the paracrine TNFα signaling (Chaudhry et al. 2013).

## Discussion

Our study used 10× genomics sequencing technology for the long-read genomic DNA sequencing of warthogs and Kenyan domestic pigs that are highly adaptable to the natural environment in Africa. The two assembled genomes of the warthog and Kenyan pig families were 2.417 Gb and 2.445 Gb, respectively. Altogether, 17,781 and 22,127 coding genes, respectively, were identified through annotation. These results will provide valuable resources and data support for future evolutionary research on African pigs and genetic research adapted to the unique local geographic environment. Compared with these published genomes, the size of the two genomes assembled in this study and the length and number of N50 scaffolds were within a reasonable range.

Phylogenetic tree analysis, principal component analysis, and population structure analysis revealed that common warthogs did not belong to the *Sus* genus and were analyzed as an outer group. Additionally, we found that Kenyan and Nigerian domestic pigs belong to the same branch of European pigs. By further combining the PSMC results, we found that the Duroc pigs and African domestic pigs overlapped in approximately 3,000 years. Additionally, our results uncovered a phenomenon of gene infiltration between southern Chinese pigs and Kenyan domestic pigs. Many studies have found that Kenyan pigs in East Africa not only have European ancestors, but also have high frequency alleles from the Far East (Amills et al. 2013; Noce et al. 2015). Approximately 3,000 years ago there was a resurgence of humans to Africa. The population of Kenya, Tanzania, and Ethiopia flooded with people from Eurasia (Vansina 1995). Perhaps it was due to this population that the genes of European pigs infiltrated African pigs. Later, in the 17^th^ century, with the Indian Ocean trade, the genes of southern Chinese pigs infiltrated African domestic pigs. The gradual colonization of Africa in modern times and the strengthening of European commercial pig breeds, resulted in the dominant genes of European pigs gradually being fixed in East Africa. Domestic pigs and some genes of native African domestic pigs have almost disappeared.

By comparing the genome structure of the warthog to the genomes of other domestic pigs, we found that the warthog completely lacks the *LDHB* gene of the reference genome located on chromosome 2. The *LDHB* gene has been shown to play an important role in the infection of classical swine fever, and its overexpression can reduce the replication of the classical swine fever virus (Fan et al. 2021). The expression levels of the *LDHB* gene in the heart, lung, liver, and spleen of warthogs were compared with those of Kenyan domestic pigs. The expression level of the *LDHB* gene in the four tissues was significantly higher in warthogs than in Kenyan domestic pigs by more than 10 times (**Fig. 4c**). To verify whether *LDHB* has an inhibitory function on ASFV, the expression of fluorescent proteins in 3D4/21 cells transfected with plasmids after infection with ASFV SY18 (**Fig. 4a and b**) was observed. The results of the overexpression and RNA interference of *LDHB* proved its inhibitory effect on the replication of ASFV.

It is worth noting that in this study we found that warthogs and wild boars in southern China had gene infiltration. However, this result has not been reported in other studies. In this study, the Chinese wild boar was used as the southern Chinese wild boar. The similarity of the climate between East Africa and southern China may have led to the convergent evolution of the two. However, other algorithms are required to verify the accuracy of the results.

Combine with our gene family evolution analysis and signature analysis, gene TRIM family genes were expanded in the Duroc and contracted in the Kenya domestic pig and Warthog. Several studies have shown that TRIM family genes were involved in the negative regulation of viral transcription (Uchil et al. 2008) and positive regulation of I-kappaB kinase/NF-kappaB signaling (Uchil et al. 2013). Previous studies reported that the proteins recoded by ASFV could inhibit the activation of NF-κB (Revilla et al. 1998). TRIM family genes may be one of the reasons that warthog and Kenya domestic pigs showed resistance to ASFV (Okoth et al. 2013; Mujibi et al. 2018).

In conclusion, in this study we assembled the genomes of warthogs and Kenyan domestic pigs and explored the evolution and gene introgression between African pigs and other pigs. At the same time, and while taking into account the transcriptomic data of warthogs and Kenyan domestic pigs, we demonstrated that *LDHB* could inhibit the replication of ASFV. These newly discovered molecular genetic markers for resistance against ASF will help improve the ASF resistance of different pig breeds.

## Methods

### Sample preparation

To understand the evolution of the African Suidae, we assembled warthog and Kenyan domestic pig genomes. Tissues including the heart, spleen, kidney, and lung used in the study were removed from a wild warthog in Nairobi National Park, Kenya. Tissues of Kenyan domestic pigs, including the kidney, heart, liver, spleen, lung, tonsil, gastro-hepatic lymph node (GHLN), mesenteric lymph node (MSLN), and submandibular lymph node (SMLN) were collected from six Kenyan domestic pigs, in Homa Bay, Kenya. DNA and RNA were isolated from these tissues and used for sequencing. The DNA isolated from the liver of two pigs was used for genome assembly, and RNAs from all the tissues mentioned above were used for genome annotation (**Supplementary Table 14**). The assembled genomes of the other 15 pigs (**Table 1**) were downloaded from NCBI Assembly (https://www.ncbi.nlm.nih.gov/assembly). A total of 14 common warthogs and seven Kenyan domestic pigs were sampled from three localities around Kenya, and six domestic pigs were collected from Nigeria. Five or six pig ears were collected from each Chinese breed, including Nanyang, Bamei, Tongcheng, Neijiang, Huai, Meishan, Bama, Diannanxiaoer, Lantang, and Mashen (**Supplementary Table 10**). The individuals mentioned above were genetically unrelated.

Animals’ work and related samples collection and treatments were approved by the Institutional Animal Care and Use Committee (IACUC) of International Livestock Research Institute (ILRI) (ref no. IACUC-RC2018-17). All procedures were conducted according to the ILRI IACUC protocol 11.

### DNA and RNA extraction and sequencing

Genomic DNA was extracted from the tissues of warthogs and Kenyan domestic pigs using the DNA MagAttract HMW DNA Kit (**Qiagen, Germantown, MD, USA**) for 10× genomics according to the manufacturer’s protocol. The 10× Chromium library was prepared according to the manufacturer’s instructions (Chromium™ Genome Library Kit & Gel Bead Kit v2) and sequenced on an Illumina HiSeq2500 with a 2×250 bp read metric.

Genomic DNA was extracted from tissues or PBMCs using DNeasy Blood & Tissue Kits (**Qiagen**) according to established protocols and checked for quality and quantity using NanoDrop2000 (Thermo Fisher Scientific, Waltham, MA, USA) and agarose gel electrophoresis. A total DNA amount of 1 μg or higher was used for DNA sequencing. Total DNA was sequenced using an Illumina HiSeq 2500 sequencing system (Compass, Beijing, China).

Total RNA was isolated from tissues according to the standard protocols of the TRIzol method (Invitrogen, Carlsbad, CA, USA). RNA degradation and contamination were monitored using 1% agarose gel electrophoresis. The concentration of total RNA was measured using the Qubit RNA Assay Kit in a Qubit 2.0 Flurometer (Life Technologies, Carlsbad, CA, United States). RNA samples that met the criteria of having an RNA integrity number (RIN) value of 7.0 or higher and a total RNA amount of 5 μg or higher were included and batched for RNA sequencing. RNA sequencing libraries were constructed using the Kapa RiboErase (Roche, Basel, Switzerland), with 3 μg of rRNA-depleted RNA, according to the manufacturer’s recommendations. Libraries were sequenced using the Illumina NovaSeq 6000 S4 platform according to the manufacturer’s instructions, with a data size per sample of a minimum of 5 G clean reads (corresponding to 150 bp paired-end reads).

The sequenced DNA-Seq and RNA-Seq raw data are available from the NCBI Sequence Read Archive with the BioProject number PRJNA691462.

### Reads Alignment to the Reference Genome Sus Scrofa 11.1

The RNA-Seq raw data were trimmed based on the quality control for downstream analyses by following steps: Firstly, BBmap (Version v0.38) automatically detected the adapter sequence of reads and removed those reads containing Illumina adapters (Brian 2014). The Q20, Q30, and GC content of the clean data were also calculated by FastQC for quality control and filtering (Andrews 2010). Secondly, the resulting sequences were mapped to the reference genome (Sus scrofa 11.1) by HISAT2 (Dobin et al. 2013). NCBI Sus scrofa11.1 annotation was used as the transcript model reference for the alignment, as well as for all protein-coding genes and isoform expression-level quantifications. Finally, FeatureCounts (from subread v2.0.1) was used to calculate the number of the read counts (Liao et al. 2014).

### Genome assembly

The genome was assembled using the default parameters of the Supernova assembler (v2) designed by Illumina 10× genomics. The scaffolds were then connected using the pseudogap style to obtain the genomic draft. Fixed gaps were added between the two scaffolds, and Sealer (Paulino et al. 2015) and Gapcloser (Xu et al. 2019; Xu et al. 2020) were used to fill the gaps.

### Evaluation of genome assembly

To evaluate genome quality, the raw reads were first mapped to the warthog and Kenyan domestic pig genomes with BWA v0.7.17 (Li and Durbin 2009). Next, the coding gene sequences and transcripts of *Sus Scrofa* 11.1 were mapped to the assembled genomes, and genome completeness was verified by 4104 benchmarking universal single-copy orthologs to the genome using BUSCO v.3.0.2b (Simao et al. 2015). Finally, two assembled genomes were mapped to *Sus Scrofa* 11.1 using MUMer v3.23 (Kurtz et al. 2004). The parameter was used with “-maxmatch -1 100 –c500.”

### The identification of presence-absence variation (PAV)

The assembled genomes, including warthog and Kenyan domestic pigs, and data from 14 additional genomes including those of Landrace, Largewhite, Pietrain, ½ Landrace* ¼ Duroc* ¼ Yorkshire, Berkshire, Hampshire, Wuzhishan, Göttingen, Bama, Jinhua, Meishan, Tibetan, Bamei, and Rongchang pigs, downloaded from public databases (Table S11), were mapped to the Duroc genome (*Sus Scrofa* 11.1) using Blastn (Altschul et al. 1990) to identify the presence-absence variation in each genome.

### Read alignment and variant calling of DNA sequence reads

To facilitate better read mapping, three criteria of quality control (QC) were carried out by FastQC with “-q 20 --thread=6 --length_required=120 --n_base_limit=6.” Filtered reads from all individuals were aligned to the *Sus scrofa* 11.1 reference genome using the Burrows-Wheeler Aligner (BWA v0.7.17). We performed duplicate marking, base quality recalibration, duplicated read removal, and mapping statistics (i.e., coverage of depth) by Picard, GATK, and SAMtools. Ultimately, the alignment files (bam) were used directly for subsequent analyses.

The BAM files of 123 pigs were used for SNV detection on a population scale using GATK (v4.1.4). The GATK command was run with the functions “HaplotypeCaller,” “GenomicsDBImport,” “GenotypeGVCFs,” “MergeVcfs,” and “SelectVariants” to generate genotype calls in Variant Call Format (VCF). Moreover, the GATK command was run with parameters “QD < 2.0, QUAL < 30.0, FS > 200.0 ReadPosRankSum < -20.0” to filter each SNV VCF file. The dbSNP database (ftp://ftp.ncbi.nih.gov/snp/organisms/archive/pig_9823/VCF/) was used to identify novel genetic variations. Gene structure annotation of SNPs was performed using ANNOVAR (Wang et al. 2010). The reference genome was *Sus scrofa* 11.1 and the annotation file was downloaded from UCSC (https://hgdownload.soe.ucsc.edu/goldenPath/susScr11).

### Annotation of assembled genomes

Genome annotation includes two main sections: structural annotation and functional annotation. Structural annotation specifically includes repetitive sequence identification, non-coding gene prediction, and coding gene prediction.

The repetitive sequence consisted of three parts: ➀ de novo prediction: repetitive sequences of the whole genome were predicted using RepeatModeler (2.0.1) (Saha et al. 2008); ➁ the repetitive sequences were mapped to the UniProt database using BLAST (Altschul et al. 1990) to remove sequences with lengths greater than 50 bp; ➃ data-driven homology annotation: the repetitive sequences obtained in the previous two steps were merged into RepBase (v20181026) (Saha et al. 2008) and then RepeatMasker (v4.1.1) (Bergman and Quesneville 2007) was used to perform soft annotation at the whole genome level.

Three procedures pertained to non-coding and coding genes: ➀ de novo prediction: AUGUSTUS (v3.4.0) (Stanke and Morgenstern 2005) and GeneMark-ES (v4.62) (Lomsadze et al. 2005) were used for genome prediction; ➁ data-driven homology annotation: HISAT2 (v2) (Kim et al. 2019) was used to perform quality control of RNA-seq data, StringTie (Pertea et al. 2015) was used to assemble the transcripts, and TransDecoder (Haas et al. 2013) was used to predict the open reading frame (ORF) structure; ➃ combining the prediction results: the results of the first two steps were filtered and merged to obtain the final annotation results using EVidenceModeler (Haas et al. 2008).

The functional annotation consisted of the following two steps: ➀ the predicted proteins were aligned with the UniProt database using DIAMOND (Buchfink et al. 2015); ➁ Pfam domain annotation was performed using PfamScan (Jones et al. 2014; El-Gebali et al. 2019) based on the Pfam database.

### Annotation of merged genomes

The *Sus Scrofa 11*.*1* genome was merged with the unique sequences of the warthog and 14 additional domestic pig genomes. Repetitive sequence prediction was ignored owing to the short sequences.

### Non-coding genes and coding gene prediction

The *Sus scrofa* of OrthoDB 10.1 (https://www.orthodb.org/) was used as the protein data set for homology comparison, and BRAKER2.1.5 (Bruna et al. 2021) software was used for gene structure prediction. In the BRAKER2 process, GeneMark-ES (Lomsadze et al. 2005) was used to perform de novo analysis of the genome to obtain seed genes. Then, the seed genes were mapped to the protein data set using DIAMOND (Buchfink et al. 2015) with fine mapping of the splicing sites using Spaln v2.3.3d (Gotoh 2008). Using the mapping information, high-quality genes of the genome were obtained using GeneMark-EP+ (Bruna et al. 2020). Finally, the high-quality genes used as the training set and the AUGUSTUS (v3.4.0) (Stanke and Morgenstern 2005) parameters were trained to obtain the final structure prediction result.

### Function annotation

Blastx (Altschul et al. 1990) was used to map the obtained protein sequences to the non-redundant protein sequence database (evalue=1e-6, minimum percentage of identical matches=70%). InterProScan (v5.48-83.0) (Jones et al. 2014) was used to annotate the protein domains and GO terms. The classification and quantity of repetitive elements were determined using the Dfam_3.1 (https://www.dfam.org/releases/Dfam_3.1/) mapping to the pig-specific library of Repbase TE using RepeatMasker with rmblastn version 2.10.0. The Kimura two-parameter model (Nishimaki and Sato 2019) was used to calculate the degree of divergence of the transposon sequence. The CpG content of the RepeatMasker sequence was used to correct the divergence level, and the correction model was DCpG = D/(1 + 9FCpG). The transposon activity time was estimated using the annual replacement rate of 5×10^−9^ (substitutions/site per year). The distribution histogram of the transposon sequence divergence was drawn using a bin size of 0.01.

### Phylogeny construction and estimate of divergence time

We constructed a phylogenetic tree of warthogs, Kenyan pigs, Duroc pigs, cattle, mice, and humans using the maximum likelihood analysis of a concatenated alignment of 3,489 single-copy orthologous genes shared with their genomes [PhyML (Guindon et al. 2010)]. iTOL was used to reconstruct and optimize the phylogenetic tree. The divergence time among the warthogs and Kenyan and Duroc pigs was estimated using the MCMCtree program as implemented in the phylogenetic analysis of the maximum likelihood (PAML) package (Yang 1997). The calibration times (89.3, 96.8, and 62.0 mya) were derived from the TimeTree database.

### Gene family expansion and contraction

We determined the expansion and contraction of the gene ortholog clusters by comparing the cluster size differences between the ancestor and each of the current warthog, Kenyan, Duroc, cattle, mouse, and human clusters (P < 0.05) using OrthoFinder (Emms and Kelly 2019) and the Café program (De Bie et al. 2006), which is based on a probabilistic graphical model.

### Phylogenetic analysis

The distance between the genomes was determined using Minhash (Berlin et al. 2015). First, genome sketches were conducted using different genome sequences to complete the calculation of different genome distance matrices and the construction of evolutionary trees. Finally, the constructed evolutionary tree was visualized using iTOL software (Letunic and Bork 2019).

The demographic history of warthogs, Kenyan domestic pigs, Duroc, Landrace, and Meishan pigs, and Chinese wild boars was analyzed using PSMC v.0.6.5 software (Schiffels and Durbin 2014). The parameter generation interval (g) was 5, and the mutation rate of each generation was set to 1.25×10^−8^.

### Analysis of genetic introgression

The genomic variation of warthogs, Kenyan pigs, Nigerian domestic pigs, Netherlands wild boars, Duroc pigs, Landrace pigs, Mashen pigs, Bamei, Nanyang, Meishan, Tongcheng, Bama, Luchuan, Wuzhishan, and Chinese wild pigs were performed for gene migration using TreeMix (Drummond et al. 2012). The warthog was analyzed as an external group. Genotype frequency data were used to construct a maximum likelihood tree to determine whether there was gene flow between the two populations. Genetic introgression events were also detected using the D-statistic (ABBA-BABA test) in Dsuite (Malinsky et al. 2021). We calculated the D statistic and Z-score, using sliding window analysis with 50 kb windows. Gene introgression is considered to be existed when D statistic ≠ 0 and Z-score > 3 in the Dmin results.

### Selection signature analysis

The composite likelihood ratio (CLR) was used to detect the selection signal in the warthog population using SweepFinder (DeGiorgio et al. 2016) with a continuous window of each 100 kb of the genome.

The Fst value between warthogs and the Netherlands wild pigs as well as warthogs and southern Chinese wild boars was detected using VCFtools (Danecek et al. 2011) with an overlapping continuous window of 500 kb and a step size of 50 kb.

### Gene functional analysis

The functional KEGG pathways (Kanehisa et al. 2021) enrichment of genes using selection signature were clustered using KOBAS3.0 (http://kobas.cbi.pku.edu.cn/kobas3/genelist/). The pathways with q < 0.05 were considered significant. We confirm that KEGG pathway map images only used in our supplementary data. According to the Kanehisa Laboratories’ policy, no permission is required when KEGG pathway map images only used in the supplementary data.

Protein-protein interaction (PPI) networks were constructed using the Search Tool for the Retrieval of Interacting Genes (STRING) online database (Szklarczyk et al. 2019), with an interaction score >0.15 set as the cut-off value.

### The expression of fluorescent proteins in 3D4/21 cells after plasmid transfection

To validate the resistance of *LDHB* to ASF, the expressions of fluorescent proteins in porcine alveolar macrophage 3D4/21 cells transfected with pEGFP-C1-LDHB and siRNA-LDHB were detected. The 3D4/21 porcine alveolar macrophages used in this study were purchased from the American Type Culture Collection (ATCC CRL-2843). The entire experiment was completed at the Institute of Military Veterinary Medicine, Academy of Military Medical Science. The ASFV strain SY18 of genotype II (GenBank accession number MH766894) used in this study was supplied by the Institute of Military Veterinary Medicine, Academy of Military Medical Science. The culture and viral titer determination of ASFV SY18 are described in a previous study (Fan et al. 2020).

The pEGFP-C1-LDHB, siRNA-LDHB, blank control pEGFP-C1, and blank control siRNA vectors were constructed by TsingKe Biological Technology (Beijing, China). The forward sequence and reverse sequence of siRNA-LDHB are 5’ - GCAAGGUUUCGCUAUCUUATTUAAGAUAGCGAAACCUUGCTT – 3’ and 5’ - GCAAGGUUUCGCUAUCUUATTUAAGAUAGCGAAACCUUGCTT – 3’. Primary alveolar 3D4/21 macrophages were seeded in 6-well plates at a concentration of 2 × 10^6^/mL and incubated at 37 °C with 5% CO_2_ in an RPMI 1640 maintenance medium. When the cell density was approximately 70–90%, the pEGFP-C1-LDHB, siRNA-LDHB, pEGFP-C1, and siRNA vectors were transfected into the cells according to the lipo3000 instructions, and incubated at 37 °C with 5% CO_2_ for 24 h. The cells were then infected with ASFV strain SY18 (MOI = 5) and were digested and harvested by centrifugation at 48 h post-infection. The cells were observed using a confocal laser scanning fluorescence microscope (Olympus LSCMFV500).

## Acknowledgements

We thank International Livestock Research Institution for offering laboratory for ASFV infection. And we also thank Chen Teng, who from the Institute of Military Veterinary Medicine, Academy of Military Medical Science for the work of overexpression of gene *LDHB*.

## Data Access

The sequenced RNA-Seq raw data of these 68 samples is available from NCBI Sequences Read Archive (BioProject number: PRJNA691462).

## Supplementary materials

### Supplementary Figures

**Fig. S1** Neighbor-joining tree constructed from SNPs using MEGA X.

**Fig. S2** PCA analysis using the SNPs of 25 pig breeds.

**Fig. S3** Structure analysis from k = 2 to 10 using ADMIXTURE.

**Fig. S4** Selection signature analysis in populations.

### Supplementary Tables

**Table S1** Summary of raw sequencing data

**Table S2** Statistic of two assembled genomes

**Table S3** Quality assessments of assembled genomes

**Table S4** Specific and annotated PCGs of Warthog comparing to reference genome

**Table S5** Statistic of repetitive elements Warthog and Kenya domestic pig genomes

**Table S6** The contracted gene family observed

**Table S7** The expanded gene family observed

**Table S8** The number and length of PAV sequences

**Table S9** Protein coding gene in PAVs of Sus scrofa 11.1

**Table S10** Statistic of whole genome sequencing of 25pig breeds

**Table S11** The D statistic results of gene introgression using Dsuite

**Table S12** Analysis of KEGG enrichment of strong selective genes in warthog

**Table S13** Number of Non-synonymous SNPs in exons

**Table S14** Summaries of RNA sequencing reads

**Table S15** The overlapping genes of pi and xp-ehh results between six comparisons

**Table S16** The overlapping genes of pi and xp-ehh results between Kenya domestic pigs and Duroc

**Table S17** The overlapping genes of pi and xp-ehh results between Kenya domestic pigs and Landrace

